# Variation in anthropogenic tolerance alters dispersal capacity of a large carnivore

**DOI:** 10.1101/2025.09.29.677867

**Authors:** K. Whitney Hansen, John J. Morgan, Luca De Alfaro, Christopher C. Wilmers, Natalia Ocampo-Peñuela

**Affiliations:** Environmental Studies Department, University of California Santa Cruz, California, USA; Ceasar Kleberg Wildlife Research Institute, Texas A&M University-Kingsville, Kingsville, TX, USA; Computer Science and Engineering Department, University of California Santa Cruz, California, USA

**Keywords:** anthropogenic tolerance, connectivity, dispersal, human disturbance, individual variation, *Puma concolor*

## Abstract

Population connectivity is essential for long-term species persistence, and a major challenge to the conservation of isolated wildlife populations globally. Understanding how individual variation in movement behavior scales up to affect population connectivity across human-dominated landscapes is essential for the development and implementation of effective conservation and management plans, particularly for wide-ranging species with potential for human-wildlife conflict. Based on the observed movement behavior of 84 pumas (*Puma concolor*) in the Santa Cruz Mountains, we characterized individual variation in tolerance towards anthropogenic presence and used EcoScape, a novel functional habitat connectivity algorithm, to predict how individual differences in behavior affect the connectivity and dispersal capacity of an at-risk carnivore population. We found that individual differences in human tolerance have a greater effect than sex-based differences on modeled ability to disperse through anthropogenic landscapes, and strongly influence connectivity. This work highlights how individual variation in animal responses to human disturbance affects dispersal capacity, and offers a novel approach to predicting the persistence of metapopulations in the context of rapid global change.

## Introduction

Habitat connectivity supports long-term species persistence by increasing functional habitat and enabling movement among patches, which maintains genetic diversity and links otherwise isolated populations (Bohonak 1999). Without the ability to make long-distance movements between isolated populations (i.e., dispersals) species are at risk of extinction (Soulé 1987; Frankel and Soulé 1981). Even with the physical capacity to disperse, doing so in human-modified landscapes can be a dangerous endeavor (Lidicker and Stenseth 1992; Blanco and Cortés 2007; Fletcher Jr. et al. 2019; Maag et al. 2019; Ash et al. 2020; Vasudev et al. 2023). Long-distance movements can bring wildlife into greater contact with humans, risking survival and thus successful dispersal (Vasudev et al. 2023). To understand and predict dispersal at the landscape scale, we need to quantify both structural (i.e. contiguous natural habitat between target habitat patches) and functional (i.e. actual movement of organisms among patches) connectivity (Kindlmann and Burel 2008).

Anthropogenic disturbances reduce landscape connectivity and limit both the likelihood and extent of animal dispersal (Tucker et al. 2018; Naude et al. 2020). In pumas living in developed landscapes (*Puma concolor*), human presence restricts available habitat needed for reproductive behaviors (Wilmers et al. 2013), and, after long-term exposure to human development, decreases survival (Nisi et al. 2023; Benson et al. 2023). To compensate for disturbance caused by human presence, pumas increase kill rates (Smith et al. 2017), increase calorie expenditure (Wang et al. 2017), and alter activity patterns (Nickel et al. 2020; Bolas et al. 2025), Dispersing individuals must face these compounded pressures in anthropogenically disturbed areas, as well as the riskiness of navigating unknown habitat (Bonte et al. 2012; Forrester et al. 2015). Unsurprisingly, pumas attempting to move through heavily fragmented landscapes have noticeably decreased dispersal distances (Stoner et al. 2013) and, even in areas where hunting them is not permitted, surprisingly low survivorship (55.8%; Vickers et al. 2015).

Certain species or individuals can be more or less tolerant of anthropogenic disturbance, which may mediate their behavioral responses to humans (Samia et al. 2015; Bouyer et al. 2015). Individual differences in movement behavior are ubiquitous across species (Hertel et al. 2020), and so individuals with varying degrees of tolerance to anthropogenic disturbance may employ different behaviors in response (Harding et al. 2019). Individuals may vary in their degree of anthropogenic tolerance due to social tendencies (Poulin 2022), sex (Wilmers et al. 2013), life stage (O’Neil et al. 2020), or intraspecific variation (Harding et al. 2019). Females with dependent young, for instance, are more sensitive to anthropogenic disturbance than males due to their needs to raise young in minimally disturbed areas (Wilmers et al. 2013; Yovovich et al. 2020), and sex-specific territoriality contributes to sex- and age-biased dispersal, with younger males more likely to disperse following the death of older territorial males (Robinson et al. 2008; Cooley et al. 2009). In a large mammalian species such as the puma, where movement and connectivity are heavily influenced by human disturbance (Tucker et al. 2018; Dickson et al. 2013), understanding how variation in tolerance to human disturbance influences dispersal capacity may be key to conservation and management (Merrick and Koprowski 2017).

Individual variation in movement behavior and dispersal may have population-level consequences (Wolf and Weissing 2012; Spiegel et al. 2017), with impacts ranging from gene flow to metapopulation dynamics (Cote et al. 2010; Spiegel et al. 2017). Indeed, differences between individuals may be a key driver of dispersal processes (Cote et al. 2010). In the western mosquitofish (*Gambusia affinis*) asocial individuals are more likely to disperse than social ones, and theoretical work has suggested that the amount of variation in sociality across individuals influences the speed of the population’s invasion front (Fogarty et al. 2011). Phenotypic differences in weasels, specifically body size, dictate dispersal capacity of individuals based on interacting effects with barriers to movement capacity, resulting in genetic structuring across habitat types (McDevitt et al. 2013). Despite the theoretical demonstrations of individual heterogeneity affecting metapopulation dynamics (Hawkes 2009), and the importance of considering individual differences in wildlife conservation strategy (Merrick and Koprowski 2017), there remains a research gap in investigating the spatial outcomes and conservation implications from individual differences in dispersal capacity of large mammals. While we know that human-wildlife conflict affects survival of dispersing individuals and, subsequently, population performance (Vasudev et al. 2023), it is less clear how individual variation in response to humans may mediate this process-specifically as it pertains to degree of anthropogenic tolerance. Here, we examine how individual differences in anthropogenic tolerance mediate the dispersal capacity and landscape connectivity of a regionally threatened large carnivore, the puma, providing insight into how behavioral heterogeneity might shape population persistence and conservation outcomes.

In California, pumas are divided into several genetically distinct populations due to habitat fragmentation. The central and south coast populations are the most isolated (Gustafson et al. 2019; 2022) and, as a result, these isolated populations are being considered for listing under the California Endangered Species Act (Yap, Cummings, and Rose 2019). Additionally, large amounts of public and private dollars are being spent to create wildlife corridors to protect land from development and facilitate highway crossings for pumas in particular (Moinuddin 2021). For a species of conservation interest such as the puma, quantifying the spatial consequences of intrinsic factors impacting movement, specifically as they pertain to degree of anthropogenic tolerance, is particularly important in management planning and spending for future connectivity. As an example, should only habituated or highly tolerant pumas be likely to disperse, then management strategies for potential human-wildlife conflict along movement corridors may be more important to design as compared to situations where less tolerant, and likely more cryptic, pumas are likely to disperse. Additionally, wildlife managers may be able to create additional corridors for those highly tolerant pumas across a broader array of suitable habitat (e.g., including areas close to human development).

To address these questions, we utilized a combination of habitat selection analyses and a novel connectivity method called EcoScape (Ocampo-Peñuela et al. 2023), with high processing capacity and a dispersion-based algorithm for simulating connectivity, to generate connectivity simulations which vary based on sex and degree of anthropogenic tolerance. We determined 1) how puma habitat selection and movement in response to anthropogenic features (e.g., degree of anthropogenic tolerance) varies between sexes and among individuals, 2) how this range in tolerance alters dispersal capacity, and 3) how and whether individuals can disperse across broader central coast region. The goal of this work is not to determine the exact level of connectivity between the Santa Cruz Mountains and adjacent mountain ranges, but rather to use it as a case study to understand how puma behavior and degree of individual variation to human development modulates connectivity.

## Results

### Individual variation in anthropogenic tolerance

We fit separate iSSF’s to 84 unique pumas. Pumas tended to select for higher vegetation cover, and against steep slopes, building densities, urban centers, and anthropogenic landcover (Figure 2). However, the anthropogenic covariates of male pumas had more overlap with 0 than females?, indicating a more relaxed avoidance, on average, to anthropogenic features on the landscape (Figure 2). Males in particular had a broad range of responses to distance to urban edge (Figure 2). However, we found that tolerance-specific differences outweighed sex-specific differences, such that high tolerance individuals across sexes had more similar habitat selection coefficient estimates than same-sex individuals of varying tolerance (Figure 2). Our averaged Spearman correlation between predicted and ideal ranks across all folds for all individual models was 0.69, indicating a moderate predictive ability of our models.

### Tolerance-specific permeability

We generated spatial predictions of habitat permeability by sex and tolerance level across the Central Coast (Figure 3). As suggested by our model results, we found stark spatial differences in landscape permeability according to tolerance level, and small differences between the sexes. Generally, medium threshold males had higher permeability across the landscapes than medium threshold females. Higher tolerance pumas (with reduced avoidance of humans) had high permeability across urban places, and mostly avoided sloped areas, while lower tolerance pumas strongly avoided all urban areas, as expected by their respective covariate combinations (Table 2). Surprisingly, high tolerance females showed increased permeability as compared to higher tolerance males, mostly due to an increased selection for cover relative to their male counterparts. High tolerance individuals appear biased towards anthropogenic features with a weak selection of natural cover-this results in an apparent reduced selection of high quality puma habitat found in parts of Big Sur, a region in Monterey County (Figure 1).

**Figure 1:**
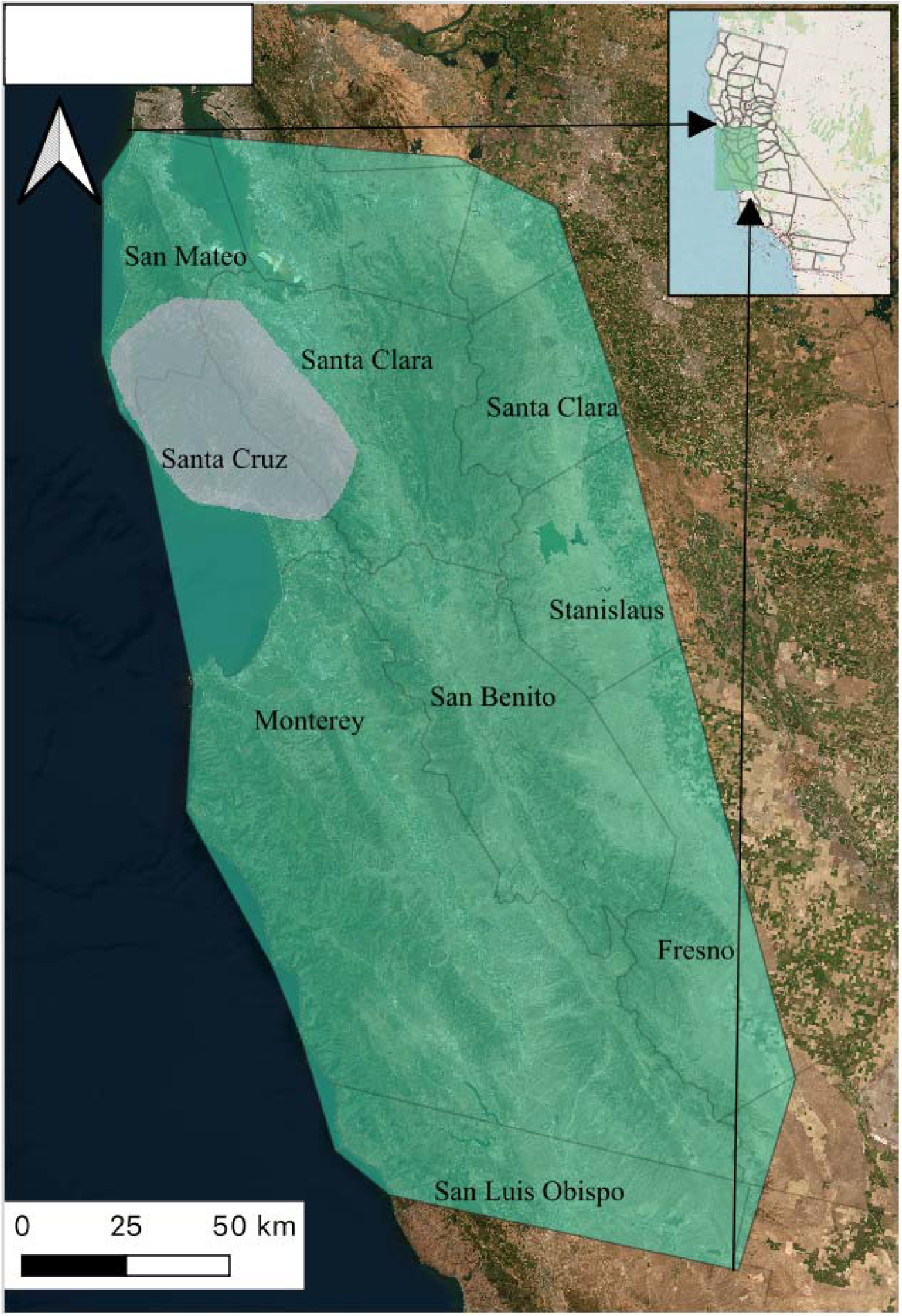
A map showing both the Santa Cruz Mountains study area (transparent white; includes parts of Santa Clara and San Mateo counties), which contained the 84 collared pumas and their historical movement data, relative to the larger region of the Central Coast (transparent teal). County lines are also included, along with labels of county names.

**Figure 2:**
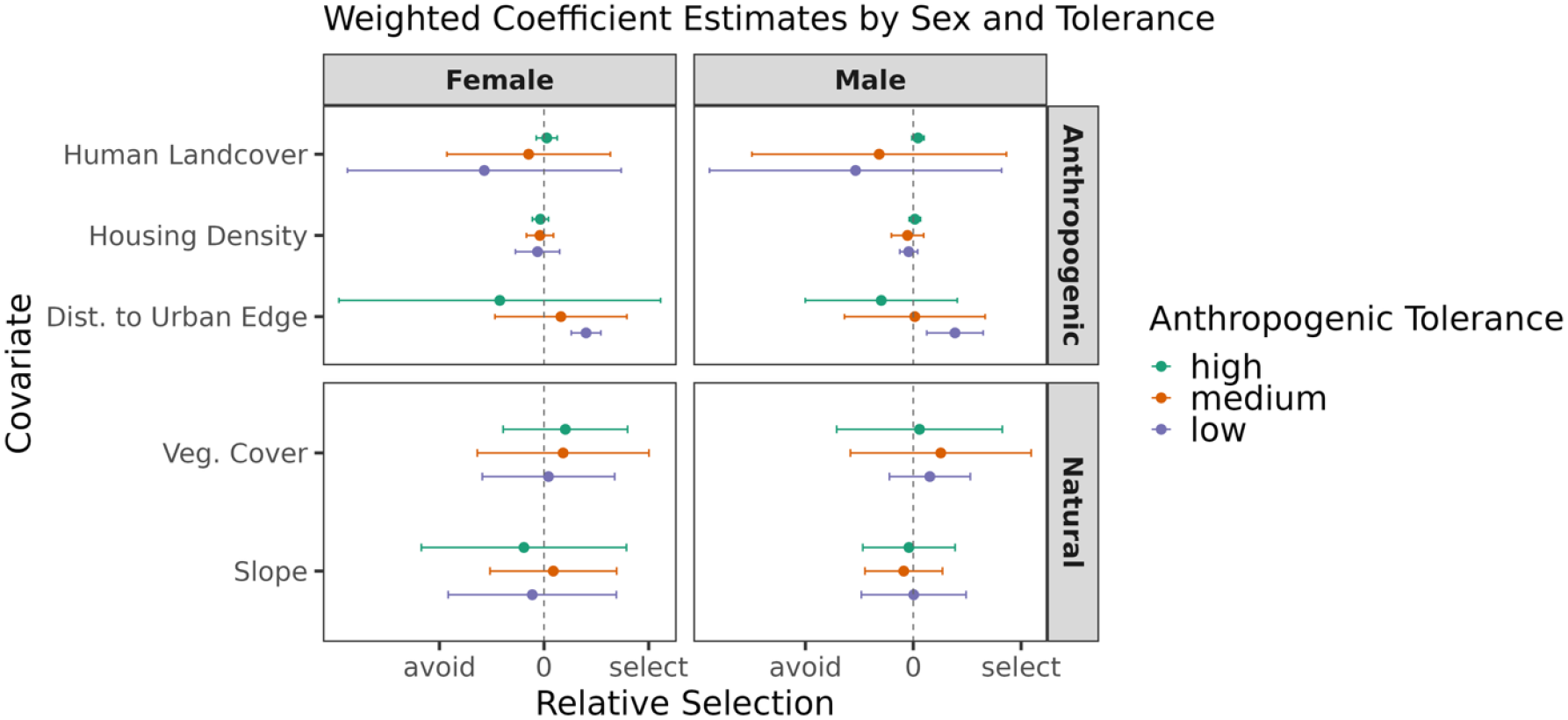
Weighted coefficient estimates for habitat features from individual iSSA’s, averaged by sex and tolerance level. Coefficient estimates are shown along a scale of relative selection to illustrate relative differences between estimates according to tolerance level.

**Figure 3:**
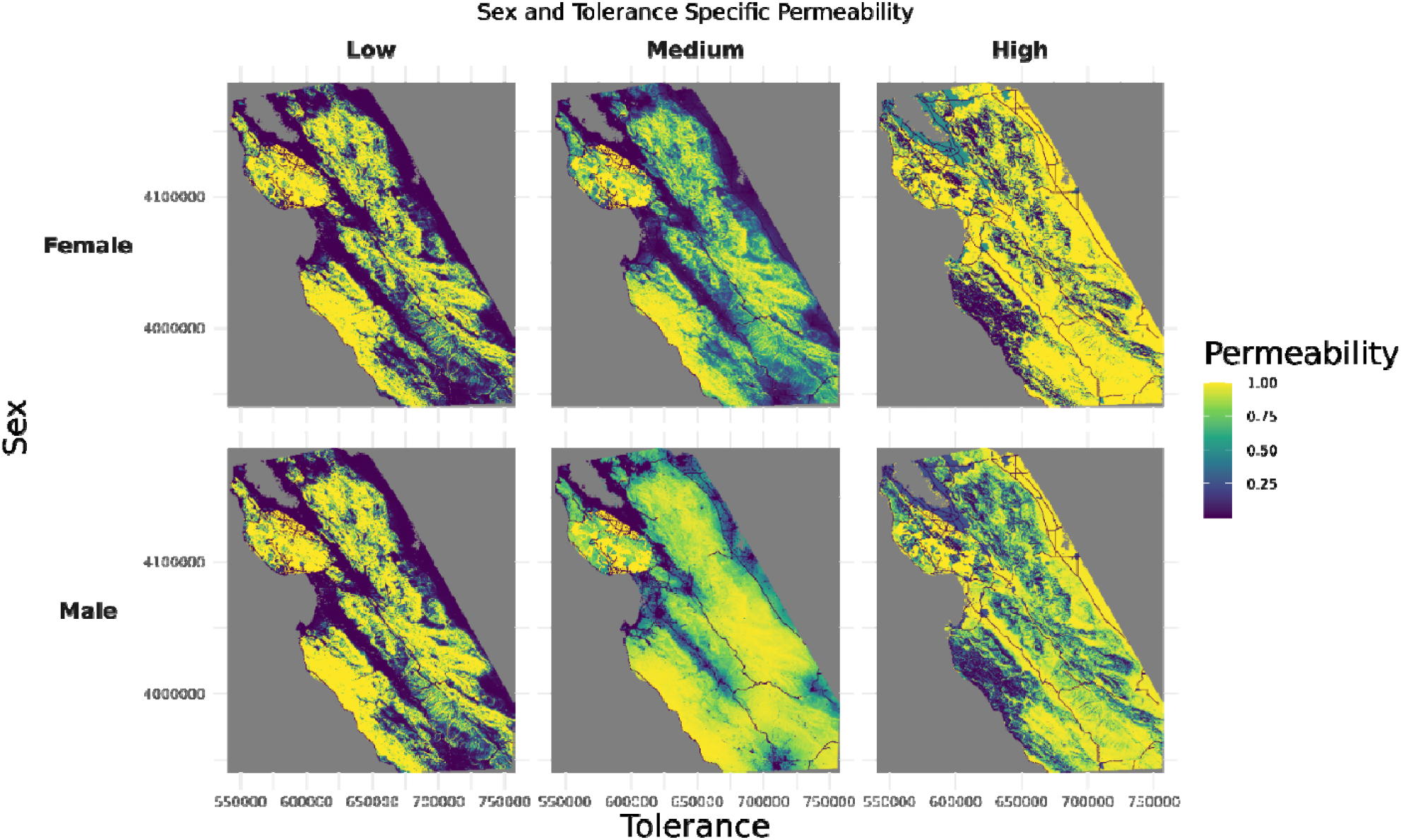
Sex and tolerance specific permeability spatial predictions. We isolated covariates from individual puma iSSA models that met specific threshold criteria for low, medium, and high tolerance per sex, and extrapolated permeability (e.g., relative selection strength) per pixel across the Central Coast of California. These maps demonstrate relative selection strength of landscape features, which means it represents preferred habitat relative to other available habitat on the landscape.

### Regional connectivity outcome

We simulated connectivity such that our theoretical pumas could start anywhere on the landscape, with a relative probability equal to the permeability of a given pixel. We found that only high tolerance individuals appeared to have consistent, region-wide dispersal capacity across both sexes, as illustrated by the lack of black-coloration subdividing Santa Clara, San Benito, and Monterey counties (Figure 4). When investigating the flow, it is clear in the low and medium threshold scenarios that both males and females are bottlenecked by unsuitable habitat (Figure 4).

**Figure 4.**
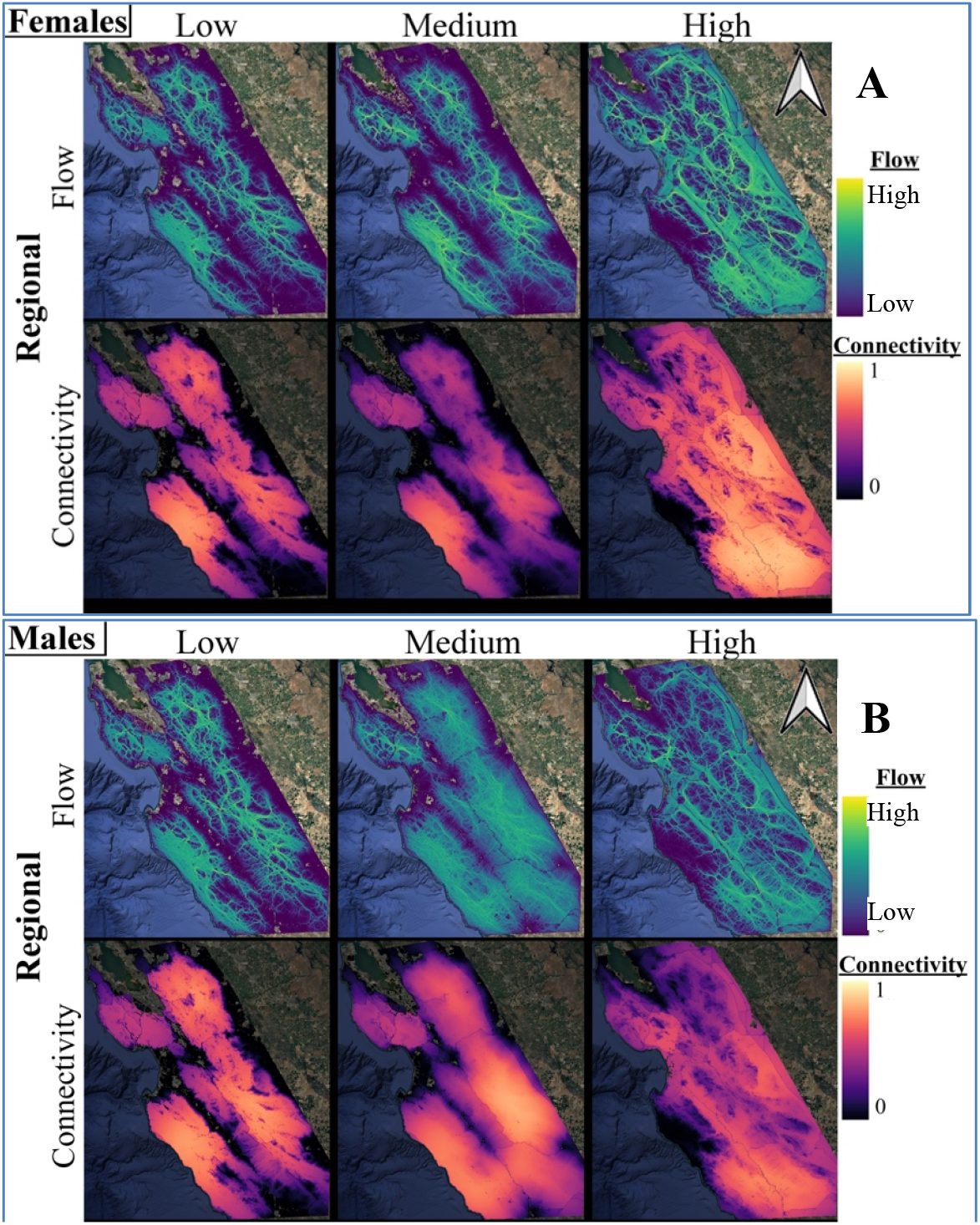
Sex and Tolerance Specific Flow and Connectivity, by Scenario. Connectivity and flow results for female (A) and male pumas (B) based on tolerance level.

## Discussion

In this study we simulated the connectivity potential of pumas across California’s Central Coast based on sex and degree of anthropogenic toleranace. While there have been numerous efforts to map and better understand puma dispersal (Beier 1995; Sweanor, Logan, and Hornocker 2000; Stoner et al. 2008; Mcrae et al. 2005; Stoner et al. 2013; Dickson et al. 2013; Ernest et al. 2014; Vickers et al. 2015; McClure, Dickson, and Nicholson 2017; Suraci, Nickel, and Wilmers 2020; Gustafson et al. 2022), and the dispersal of large carnivores in general (Crooks et al. 2011), this is the first study to do so while explicitly considering inherent, and highly relevant (Čapkun-Huot et al. 2023), variation amongst individuals that affects movement capacity in anthropogenic landscapes (see research on weasels (McDevitt et al. 2013) and birds (Doerr and Doerr 2005)). The degree of anthropogenic tolerance clearly affected the dispersal capacity of individuals. Males overall were able to achieve greater connectivity when comparing sex-specific medium threshold individuals, but the sex-specific differences were far less significant than tolerance-specific differences. Furthermore, the flow layers reveal that for low and medium tolerance individuals connectivity can only be achieved via specific bottlenecks that allow flow, many of which overlap busy highways and put the pumas at risk of vehicle collisions. This reality resulted in low probabilities of connectivity in the pixels surrounding our study area. While the population in Santa Cruz is relatively stable, the population cannot function if management strategies do not allow dispersers into the region as the population will become more inbred and lose viability (Nisi et al. 2023).

While we expected our simulations of highly tolerant pumas to suggest dispersal capacity across busy highways or urbanized areas, these spatial predictions do not account for increased expected mortalities in these same places, such as from traffic collisions (Nisi, Benson, and Wilmers 2022). Other work in California has determined that vehicle collisions are the second highest perpetrator of puma mortality events (Benson et al. 2023), after management killing. In our dataset alone, we isolated the 16 high tolerance pumas (n = 9 males, n = 7 females) to determine their fates. Of these individuals 60% have died while collared, and 50% of them died from anthropogenic causes. It is well documented in the Santa Cruz mountains that exposure to anthropogenic features increases mortality risk (Nisi et al. 2023), and so dispersing individuals that face increased exposure to anthropogenic features while attempting to navigate fragmented habitat certainly have increased mortality risk (Newby et al. 2013; Vickers et al. 2015).

Therefore, while the simulated connectivity maps for higher tolerance individuals reveals high dispersal capacity vis-a-vie movement and habitat selection, it does not reflect the realities of mortality risk en-route (Ash et al. 2020; Vasudev et al. 2023). More importantly, our results and anecdotal survival rates of highly tolerant pumas point to a more critical conservation concern, which is a potential mismatch between dispersal capacity and mortality risk. Specifically, it would be crucial to understand whether those pumas with increased tolerance and dispersal capacity through anthropogenic landscapes are actually the most at risk of anthropogenic-caused mortality, essentially nullifying the genetic potential of their dispersal capacity. Indeed, carnivores that increase exposure to humans, such as using them as shields to escape apex predators or due to learned habituation from their mothers, face increased mortality risks (Prugh et al. 2023; Shimozuru et al. 2020). Mismatches between perceived risk and the lethality of human presence may have population level consequences, and requires further investigation (Smith et al. 2024). Given the relevance of ecological traps in our system (Nisi et al. 2023), management strategies for pumas may require balancing both the spatial needs of less anthropogenically-tolerant pumas with hazing or coexistence strategies for the more tolerant pumas that increase exposure to anthropogenic presence.

Importantly, our simulations suggest low and medium tolerance individuals in both sexes, which represent 80% of the pumas in our sample, struggle to disperse across the Central Coast.

While individual variation in movement is being increasingly investigated (Hertel et al. 2020) including as a driver of population-level processes, its role in dispersal processes is less understood, particularly in mammals (Hawkes 2009). Should dispersal processes be maintained by individual variation, as suggested in some species (Duckworth 2008), then it would be pertinent to consider how the differences between dispersing individuals interact with anthropogenic presence. If only the aggressive and bolder individuals disperse (Cote et al. 2010), then coexistence strategies for contentious species, such as large carnivores, must consider connectivity planning that incorporates a spatial understanding of how these types of individuals move across the landscape (Merrick and Koprowski 2017). While we do not explicitly consider personality types or behavioral syndromes in our study (Dingemanse and Wright 2020), the degree of anthropogenic tolerance is a highly relevant and potentially straightforward trait that varies amongst individuals, and has important conservation-related consequences (Bouyer et al. 2015; Čapkun-Huot et al. 2023). Movement ecologists have characterized the functional responses of wildlife to varying available habitat (Holbrook et al. 2019), but more work is needed to link this variation to larger spatial and demographic processes (Matthiopoulos et al. 2011; 2019). Indeed, there could be opposing selective forces on wildlife with forced interactions with humans that would most benefit their survival; while individuals who avoid humans have improved survival, animals with greater tendences for anthropogenic habitat selection are more likely to disperse and propagate. The combination of avoidant individuals but anthropogenically tolerant dispersers may be required to maintain gene flow and diversity in a species such as the puma. It is clear that how wildlife responds to human disturbance affects population persistence, and capturing the range of individual responses allows a better understanding of species adaptability (Merrick and Koprowski 2017; Forsman and Wennersten 2016). Our study used a connectivity tool to illustrate how individual variation not only exceeds sex-specific differences, but has important spatial consequences that affect population persistence for an endangered species.

Wildlife managers concerned with population persistence must increasingly contend with multiple pressures of human disturbance on the landscape that have compounding, negative effects on wildlife (Abrahms et al. 2023). Human recreation, for instance, significantly alters the functional use of habitat for wildlife (Bolas et al. 2025; Morgan et al. *in review*). It is such a significant influence that failing to include it in habitat selection studies might result in connectivity corridor predictions that are inviable and unrealistic (Corradini et al. 2021). How humans utilize the landscape, and their sociopolitical context, may add greater variation in functional connectivity for wildlife (Behr et al. 2017; Ghoddousi et al. 2021). Human-wildlife conflict is another important form of functional human presence that has been rarely incorporated into connectivity models (Vasudev et al. 2023), yet is highly relevant to carnivore conservation (Karanth and Chellam 2009) and for pumas in particular (Benson et al. 2023). Moreover, our study does not consider additional stressors from climate change, which will not only increase the frequency and severity of human-wildlife conflict (Abrahms et al. 2023) but also affect the dispersal capacity of animals (McCauley and Mabry 2011) by limiting suitable habitat (Pacifici et al. 2015; Alston et al. 2020) and pushing physiological limits (Woodroffe, Groom, and McNutt 2017). In California in particular, the combinations of human population growth, the expansion of the wildland-urban interface, and climate change act synergistically to increase fire severity and extent, further modifying landscapes and limiting suitable habitat for wildlife (Lenihan et al. 2008; Hurteau et al. 2014). Connectivity planning in general needs a more integrated approach that can incorporate these stressors that compound over time (Jennings, Zeller, and Lewison 2021).

Efficient connectivity tools offer a useful way to investigate ecological questions, but also more effective conservation planning by 1) investigating connectivity over broader spatial scales, 2) incorporating dynamic elements that require multiple scenarios and future projections, and 3) investigating multiple species at once (Ocampo-Peñuela et al. 2023). We informed the EcoScape algorithm using movement and spatial data, deriving our permeability layers using well-developed movement ecology methods (Zeller et al. 2016), and GPS collar data to set algorithm parameters. In our simulation, dispersal distance limits the physical distance pumas can disperse and therefore would restrict overall connectivity. However, the variation in dispersal capacity between tolerance levels suggests that the dispersal distance parameter also functions as a way to capture an individual’s capacity to anticipate high quality habitat beyond immediate surroundings. Using EcoScape, by increasing dispersal distance indefinitely pumas will eventually reach any destination, given enough theoretical pumas are generated over enough simulations. Low and medium tolerance pumas were so restricted by barriers around the Santa Cruz study area that they were essentially unwilling to push through fragmented habitat, despite the physical ability to do so (vis-a-vie the dispersal parameter), as evidence by the low connectivity pixels surrounding the Santa Cruz area.

We present a case study that incorporates disturbance tolerance into mammal connectivity modelling to both investigate behavioral mechanisms driving individual variation in habitat selection and guide conservation decisions. This example shows the potential for EcoScape to incorporate intra-specific variability into connectivity models, unlocking the potential for more ecologically-sound and realistic movement models. We hope this inspires other researchers with movement data to use it to predict movement and guide conservation decisions across continental scales.

## Methods

### (1) Overview

We simulated three different theoreitcal puma types to test how variation in anthropogenic tolerance would affect connectivity and dispersal capacity. We simulated a movement scenario pumas can start their dispersal from anywhere in their Central Coast habitat, and the resulting map depicts overall movement capacity for dispersers across the region. We compared connectivity (probability of moving through a given landscape pixel) and flow (the relative importance of that pixel to overall connecticity) across sex and human tolerance level. Lastly, we use our findings to discuss the ramifications of individual variation in dispersal capacity, and the importance of considering intrinsic factors affecting movement in the conservation and management of wildlife populations.

We leveraged a long-term movement dataset of 84 individual pumas from the Santa Cruz Mountains to investigate habitat selection and movement. Utilizing habitat selection covariates estimated in movement models such as the integrated step selection function (iSSF) has been effective in calibrating resistance surfaces used for wildlife connectivity modeling (Zeller et al. 2018). We therefore fit an iSSF to each individual puma to quantify movement and habitat selection relative to anthropogenic presence, while controlling for natural landscape features. We classified individual pumas into degrees of anthropogenic tolerance based on observed, individual puma responses to anthropogenic landscape features. We aggregated iSSF covariates by degree of anthropogenic tolerance to generate tolerance-specific predictive surfaces: instead of inverting them (per Zeller et al. 2018) into resistance surfaces, we multiply model coefficients across available habitat to generate permeability surfaces. We then input these permeability surfaces into EcoScape to predict tolerance- and sex-specific connectivity (the probability an animal can move through an area) and flow (the importance of that area to overall connectivity).

### (2) Study area and GPS data

We used GPS data collected from free-ranging pumas captured and collared in the Santa Cruz Mountains. The Santa Cruz Mountains are in the Central Coast of California, and the 2800 km2 study area is characterized by a gradient of human development ranging from agricultural to rural to highly urban. Habitat types vary between redwood, mixed oak, and chaparral forests, as well as open grasslands and coastal shrublands. For more details, see Wilmers et al. 2013.

Pumas were captured and fitted with GPS collars (Vectronics Aerospace GPS Plus or Vertex) set to collect data at either 5 minute or 4 hour intervals. Pumas were captured using cage traps or trained hounds, following protocols approved by California Department of Fish and Wildlife (Scientific Collecting Permit ID: S-190700003-19070-001) and the University of California, Santa Cruz’s Institutional Animal Care and Use Committee (proposal code: Wilmc2212dn). Collar data in this study was aggregated from 84 individual pumas (41 females and 43 males), with data ranging from May 2009 through January 2024, excluding data collected during COVID lockdowns which represented atypical puma responses to anthropogenic presence (March 17th 2021 to May 17th 2021; Wilmers et al. 2021). We collected locations every 4 hours using Iridium satellite or direct download. We then cleaned and regularized the GPS data to a fix every four hours in RStudio, which was both an appropriate temporal scale for our research questions (Nisi, Suraci et al. 2022) and maximized the number of individuals we could include in our study. On average, pumas had 1143 relocations (min = 28, max = 7644).

### (3) Environmental data and processing

Our study explores the consequences of individual variation in anthropogenic tolerance for movement behavior and the resulting implications for population connectivity. We therefore collected a suite of environmental spatial layers which contained information relevant for puma movement and space use, including slope, tree and shrub cover, and multiple forms of anthropogenic presence, specifically distance to urban edge, housing density, and land cover type. These spatial data were selected based on previous publications in our study area and other research on coastal pumas in the state of California, which suggested the importance of these environmental features in puma habitat selection (Wilmers et al. 2013; Zeller et al. 2016; Suraci et al. 2020; Nisi, Benson, et al. 2022; Nisi, Suraci, et al. 2022). We simplified our land cover data to distinguish any given pixel between natural land cover (e.g., protected, exurban, parks) and anthropogenic land cover (e.g. suburban, urban, industrial, and agricultural landscapes). In order to estimate housing density, we utilized Microsoft’s building footprint layer (Microsoft 2018), and calculated the kernel density estimate of all buildings within 150 meters at a 30 meter pixel resolution (Wilmers et al. 2013).

### (4) Develop Permeability Layer from GPS data

We used the iSSF framework to calibrate permeability surfaces to input into EcoScape (Zeller et al. 2016). We trained our movement models on the collar data in the Santa Cruz Mountains and used their output to create permeability surfaces across the larger region to make inferences on the connectivity and dispersal capacity of individual pumas along the Central Coast of California (Figure 1). To do so, we fit an iSSF to each individual puma to measure movement in response to and habitat selection for anthropogenic presence, while controlling for natural landscape features. The iSSF jointly models habitat selection and movement from GPS data, allowing for unbiased estimation of habitation selection (Avgar et al. 2016) and dual inference on both processes. To prepare our movement data, we first excluded all steps where pumas moved less than 200 meters (Zeller et al. 2016), which functionally restricted our movement data to a majority of steps taken during the nighttime and crepuscular hours (Suraci, Nickel, and Wilmers 2020). Then we generated 20 control locations from each previously used location to simulate places pumas could have chosen to go. We sampled distances for our control points from a theoretical Gamma distribution, parameterized by our empirical movement data, and sampled directions from a uniform turning angle distribution (Avgar et al. 2016). Each used point was matched with its 20 control points in a single strata. We extracted the environmental and anthropogenic covariates at all used and control points. We standardized all covariate data by subtracting each value from its distribution’s average and dividing by the standard deviation. We tested for correlation between covariates at control points, including only those covariates with a Pearson’s correlation coefficient less than 0.7 (Hinkle et al. 2003). We then modeled the differences in spatial characteristics between the used and control points per strata using a conditional logistic regression (Fortin et al. 2005). Given that we sought to understand ranges in individual responses to habitat and landscape features, we fit 84 models (one per individual), allowing each individual to respond differently to each covariate.

Previous work in our study area has identified slope, vegetative cover, urban edge, and housing density as relevant variables dictating puma movement in the Central Coast (Nisi, Benson, et al. 2022; Nisi, Suraci, et al. 2022; Nickel et al. 2020; Wilmers et al. 2013). We therefore fit an iSSF model to each puma which estimated their response to basic environmental and anthropogenic features available across a large swatch of the Central Coast. We also included a land cover covariate (natural vs. anthropogenic land cover) to explicitly highlight agricultural areas, which affect puma behavior (Nisi et al. 2022; version 1.3.2; U.S. EPA, 2010). Lastly, we used a k-fold cross validation procedure (k = 10 folds) on each individual model to determine model robustness and out of sample predictive power (Fortin et al. 2005).

We then used our habitat selection covariates from each puma to generate ranges of theoretical puma resistances from low anthropogenic tolerance to high; we did so by quantifying thresholds of tolerance. Previous research has demonstrated sex-specific differences in both mortality risk and behavioral response to human development (Wilmers et al. 2013; Benson et al. 2023), therefore we aggregated our covariates by sex. We then isolated those covariates that were both relevant anthropogenic disturbance and showed high degrees of variation (distance to urban edge and selection for landcover; see Figure 2). We defined two covariates, distance to urban edge (*U*) and anthropogenic landcover (*H*). For each individual *i*, we extracted the coefficient estimate *β*_*i,k*_ and its standard error *SE*_*i,k*_ from that individual’s iSSF. We used inverse-variance weights given by,

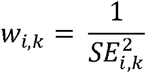

normalized to sum to one within sex

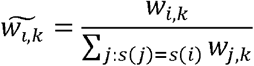

Using these normalized weights, we computed sex-specific weighted quantiles of the coefficient distributions. The weighted quantile 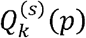 for covariate *k*, sex *s*, and quantile level *p* is given by,

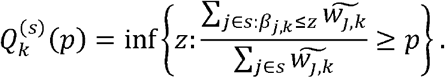

Individual tolerance categories were then assigned based on the joint position of *β*_*i,U*_ and *β*_*i,H*_ relative to these quantiles (Figure S1):

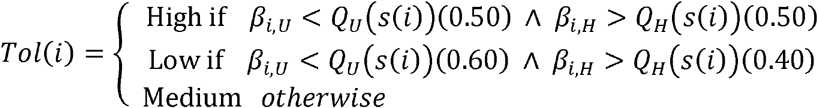

These thresholds resulted in regions of tolerance based on the relative coefficient estimate magnitude per covariate (Figure 2; Figure S1). Pumas with consistent selection for anthropogenic features were defined as high tolerance, those with consistent avoidance were defined as low tolerance, and pumas with either medium selection or a mixed strategy (e.g., selection of one feature but avoidance of the other) were defined as medium tolerance.

### (5) Theoretical pumas

We used these weighted statistical thresholds to generate three “theoretical pumas” per sex, ranging from low, medium, to high anthropogenic tolerance, by averaging all covariates (including those for cover, slope, and housing density) across the pumas grouped by tolerance level to generate a single representative puma. To validate our selection of thresholds, we considered the relationship between the two covariates in question (distance to urban edge and human landcover) along a two-dimensional graph. We hypothesized that high to low tolerance individuals would fall along a line with a −1:1 relationship between selection for distance to urban edge and selection for landcover, respectively, such that low tolerance individuals would choose to be far from urban edge and avoid landcover, while high tolerance individuals who choose the opposite (Figure S1). Medium tolerance individuals may avoid urban edge, but have relaxed avoidance of landacover (or vice versa). We modelled the linear relationship between the two covariates, per sex, and visually inspected that the theoretical pumas’ coefficient values fell, evenly spaced, along this line (Figure S1).

We used environmental data spanning the Central Coast (see Figure 1) to create a new dataset, upon which we projected relative habitat selection of pumas. In order to maintain the same degree of scaling across our broader region of California, we used the same z-score standardization procedure as before but used the mean and standard deviation values used to scale the environmental data used for each puma’s iSSA (e.g., restricted to the Santa Cruz mountains) (e.g., Hofmann et al. 2023). Functionally, this meant we were measuring selection relative to habitat available in the study area where the GPS data was collected. We set movement covariates to their means (0) to standardize movement capacity across our spatial predictions. We generated predictions for relative habitat selection across the region per puma based on the tolerance-specific combination of coefficients, resulting in a permeability raster per sex per degree of anthropogenic tolerance (n = 6). Lastly, we fit masks to each permeability raster to force our spatial prediction to represent biological realities of puma dispersal capacity in the following ways: first, all protected areas in the Santa Cruz area were given a permeability value of 1 to represent “source” populations (Nisi et al. 2023); second, we took a roads layer of all public roads in the region and assigned it a weight based on its scaled average daily traffic value. Specifically, we used a log-inverse transformation to compress the dynamic range of traffic values, and reduce the disproportionate influence of high-traffic pixels, such as 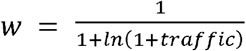. We applied these weights over our permeability layers to shrink a pixel’s relative permeability which are meant to account for the mortality risk from crossing roads (Benson et al. 2023).

### (6) EcoScape and Dispersal Assumptions

To generate our connectivity layers, we fed our permeability rasters into EcoScape (Ocampo-Peñuela et al. 2023), an efficient, propagation-based connectivity algorithm informed by metapopulation theory. EcoScape uses the values from permeability rasters to determine the propagation capacity of theoretical “seeds” (e.g. pumas) across habitat pixels on the landscape, which it runs over multiple iterations to generate both a connectivity layer (the probability that a pixel is repopulated in all simulations) and a flow layer (the importance of each pixel to overall connectivity) (Ocampo-Peñuela et al. 2023). Seeds in EcoScape can propagate up to a predetermined distance, a parameter the user sets which is relative to the pixel width of the underlying permeability raster. It is not a population model in which the seeds multiply or reproduce. The process of propagation is essentially a movement simulation where seeds are generated anywhere within predetermined habitat areas, set by the user, and then are allowed to move step-by-step through the landscape’s pixel matrix, with their movement influenced by parameters set by the user and the permeability raster. Seeds therefore preferentially move through suitable habitat and face resistance in areas they avoid (e.g., in our study, the spatial output from an iSSA which quantifies preferred habitat). The density at which seeds are generated can be a constant value, which allows a consistent amount of theoretical animals to move from any given starting location. Given that seeds can “bump” into each other, density can also be scaled to account for a more restricted starting area, such that seeds are generated at a density relative to pixel area. The propagation process spreads the seeds outward, essentially mapping probable movement routes. The biological assumptions around dispersal in a given simulation depends on the parameterization of the model, which we describe below.

Apart from the underlying permeability landscape, there are three principal parameters required to run EcoScape that are relevant to fine tuning the model for mammals and for specific scenarios: 1^st^ is how far the seed can travel (e.g. dispersal distance), 2^nd^ is the density at which seeds are generated on the landscape, and 3rd is identifying the habitat pixels from which repopulation can start (e.g. pixels where seeds can be generated). We set parameters to simulate overall dispersal capacity across the Central Coast. We set dispersal distance using our movement data. We determined this distance by identifying dispersing individuals (by visualizing data in ArcGIS), and calculated the largest distance between any two GPS points within each dispersing individual’s movement track. We selected the largest overall dispersal distance (across all dispersing individuals) as our dispersal distance parameter. While the average dispersal distance of pumas in our dataset was 44.8km, which is fairly similar to observed dispersal distances seen in other fragmented landscapes (Stoner et al. 2013), one puma in our dataset of 84 individuals dispersed 89km. In order to set a realized dispersal distance, we chose this maximum value of 89km as our parameter. We set our seed density to 4, which generated a constant number of theoretical pumas across the entire landscape. In this scenario, the probability a seed was generated on a pixel in the landscape was directly related to the pixel’s relative selection strength (Avgar et al. 2017; Zeller et al. 2016). While this resulted in pumas appearing in less than ideal habitat (albeit with a lower probability), it allowed pumas to begin dispersals from any place in the Central Coast. We generated 6 connectivity and 6 flow layers (n = 3 per sex per connectivity/flow).

## Acknowledgements

We thank Richie King, Paul Houghtaling, Cliff Wylie, Dan Tichenor, and the rest of the Santa Cruz Puma Project Team for their assistance in collecting mountain lion movement data over the years. We thank the California Futures: Scenario-Based Support for Climate-Resilient Conservation Decisions research team for their support and contributions to this project.

## Author contributions

KWH and NOP conceived the study. KWH performed the analysis, writing, the interpretation. JJM and LDA assisted with analysis, writing, and interpretation. NOP co-led interpretation, edited the manuscript, and secured funding for the research. CCW led data collection efforts, edited the manuscript and assisted in interpretation. All authors contributed to its revision and final preparation.

## Data Availability Statement

All data associated with this project are available to reviewers on Dryad at the following private link: https://doi.org/10.5061/dryad.44j0zpctn

All data will be made publicly available after review.

Associated R code is available on GitHub (https://github.com/WhitneyH1317/puma_permeability_calibration)

## Additional Information

The authors have no competing interests to declare. Pip install ecoscape_connectivity==0.2.5

