## Supplemental Figure 1 for "Variation in anthropogenic tolerance alters dispersal capacity of a large carnivore"

**Supplementary Materials:**

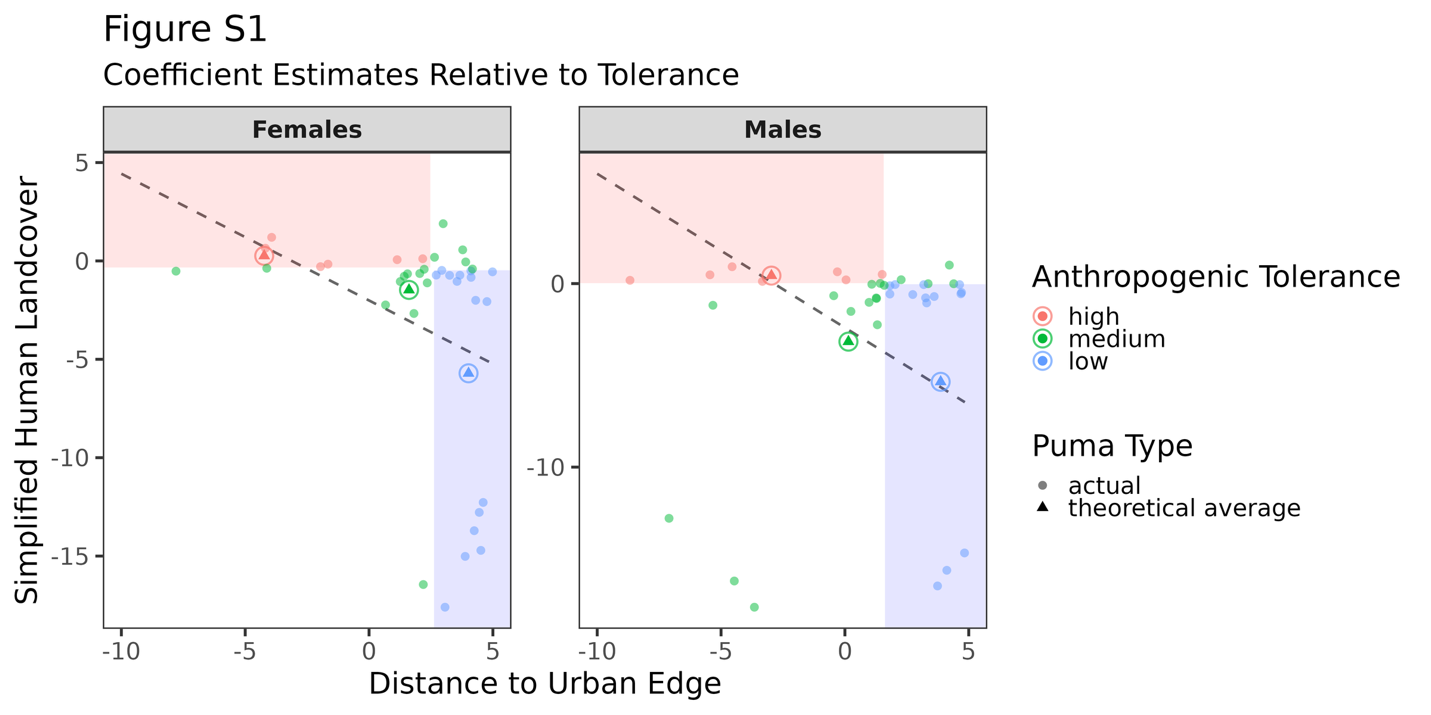

Figure S1: Coefficient estimates for human landcover (y-axis) and distance to urban edge (x-axis) per individual puma by sex. We colored in regions depicting high or low tolerance in red and blue, respectively, based on identified thresholds used to group individuals according to tolerance level. We also modelled the relationship between coefficient estimates per sex using a generalized linear model, and show the linear relationship on the plot as a dotted black line. The theoretical averaged pumas, whose respective coefficients are the result of tolerance-specific averages, are shown as circled triangles. Here we illustrate how the theoretical averaged fall relatively close to the negative linear relationship between the two covariates, illustrating a continuous range of tolerance levels among individual pumas by sex.
